# A systematic genetic analysis and visualization of phenotypic heterogeneity among orofacial cleft GWAS signals

**DOI:** 10.1101/332270

**Authors:** Jenna C. Carlson, Deepti Anand, Azeez Butali, Carmen J. Buxo, Kaare Christensen, Frederic Deleyiannis, Jacqueline T. Hecht, Lina M. Moreno, Ieda M. Orioli, Carmencita Padilla, John R. Shaffer, Alexandre R. Vieira, George L. Wehby, Seth M. Weinberg, Jeffrey C. Murray, Terri H. Beaty, Irfan Saadi, Salil A. Lachke, Mary L. Marazita, Eleanor Feingold, Elizabeth J. Leslie

**Affiliations:** Department of Biostatistics, Graduate School of Public Health, University of Pittsburgh, Pittsburgh, PA, 15261, USA; Department of Human Genetics, Graduate School of Public Health, University of Pittsburgh, Pittsburgh, PA, 15261, USA; Department of Biological Sciences, University of Delaware, Newark, DE 19716 USA; Department of Oral Pathology, Radiology and Medicine, University of Iowa, IA 52242, USA; Dental and Craniofacial Genomics Core, School of Dental Medicine, University of Puerto Rico, San Juan, Puerto Rico, 00936, USA; Department of Epidemiology, Institute of Public Health, University of Southern Denmark, Odense, DK-5230, Denmark; UCHealth Plastic and Reconstructive Surgery, Colorado Springs, CO, 80907, USA; Department of Pediatrics, McGovern Medical School and School of Dentistry UT Health at Houston, Houston, TX, 77030, USA; Department of Orthodontics, College of Dentistry, University of Iowa, Iowa City, IA, 52242, USA; INAGEMP (National Institute of Population Medical Genetics), Porto Alegre, 91501-970, Brazil; ECLAMC (Latin American Collaborative Study of Congenital Malformations) at Department of Genetics, Federal University of Rio de Janeiro, Rio de Janeiro, 21941-902, Brazil; Department of Pediatrics, College of Medicine; Institute of Human Genetics, National Institutes of Health; University of the Philippines Manila, Manila, 1000, The Philippines; Philippine Genome Center, University of the Philippines System, Quezon City, 1101, The Philippines; Department of Oral Biology, School of Dental Medicine, University of Pittsburgh, Pittsburgh PA,15219, USA; Department of Health Management and Policy, College of Public Health, University of Iowa, Iowa City, IA, 52242, USA; Center for Craniofacial and Dental Genetics, School of Dental Medicine, and Clinical and Translational Science, School of Medicine, University of Pittsburgh, Pittsburgh, PA, 15219, USA; Department of Pediatrics, Carver College of Medicine, University of Iowa, Iowa City, Iowa,52242, USA; Department of Epidemiology, Johns Hopkins Bloomberg School of Public Health, Baltimore MD, 21205, USA; Department of Anatomy and Cell Biology, University of Kansas Medical Center, Kansas City, KS 66160 USA; Center for Bioinformatics and Computational Biology, University of Delaware, Newark, DE 19716 USA; Department of Human Genetics, Emory University School of Medicine, Emory University, Atlanta, GA, 30322, USA

## Abstract

Phenotypic heterogeneity is a hallmark of complex traits, and genetic studies of such traits may focus on them as a single diagnostic entity or by analyzing specific components. For example, in orofacial clefting (OFC), three subtypes – cleft lip (CL), cleft lip and palate (CLP), and cleft palate (CP) have been studied separately and in combination. To further dissect the genetic architecture of OFCs and how a given associated locus may be contributing to distinct subtypes of a trait we developed a framework for quantifying and interpreting evidence of subtype-specific or shared genetic effects in complex traits. We applied this technique to create a “cleft map” of the association of 30 genetic loci with three OFC subtypes. In addition to new associations, we found loci with subtype-specific effects (e.g., *GRHL3* (CP), *WNT5A* (CLP)), as well as loci associated with two or all three subtypes. We cross-referenced these results with mouse craniofacial gene expression datasets, which identified additional promising candidate genes. However, we found no strong correlation between OFC subtypes and expression patterns. In aggregate, the cleft map revealed that neither subtype-specific nor shared genetic effects operate in isolation in OFC architecture. Our approach can be easily applied to any complex trait with distinct phenotypic subgroups.

## Introduction

Most complex human traits (defined as those with both genetic and non-genetic risk factors) exhibit some phenotypic heterogeneity and variable expression with potentially hundreds of significantly associated genetic risk factors showing strong evidence of association (i.e. achieving replicated genome-wide significance in large studies). Determining the relevance of any particular genetic risk factor at the individual or family level is a significant challenge. There are several methods to compare correlated quantitative phenotypes, including those using a multivariate regression framework and the correlation between multiple phenotypes in cohorts or samples of cases and controls. However, these methods estimate statistical correlation to measure phenotype relatedness and are not suitable for examining mutually exclusive disease subtypes. Further, these methods make comparisons based on the entire genome, which lacks the specificity of a more localized approach and could obscure biologically meaningful relationships.

We propose a targeted approach based on summary statistics from genome-wide association to sort out phenotypic heterogeneity. To illustrate this approach, we apply it to orofacial clefts (OFCs), congenital birth defects affecting the face and oral cavity. OFCs are the most common human craniofacial birth defect and combined they occur in approximately 1 in 800 live births worldwide (Leslie & Marazita, 2013). Although there are many types of OFCs, the term is most commonly used to refer to clefts of the upper lip and/or palate. For our purposes, OFCs will refer to cleft lip (CL), cleft lip with cleft palate (CLP), or cleft palate (CP), the three most common types of OFCs. There is an additional combined category of CL with or without CP (CL/P), historically felt to be distinct from CP alone due to the separate embryological origins of the upper lip and secondary palate. Thus, within the CL/P group, CL and CLP have been considered variants of the same defect that only differed in severity (Marazita, 2012).

Notably, more recent extensive epidemiological, genetic, and biological data suggest a more complex relationship between CL, CLP, and CP with both common and unique etiologic factors. In population-based studies in Denmark (Grosen et al., 2010) and Norway (Sivertsen et al., 2008), the recurrence risk for siblings was not uniform. The recurrence risks are consistently highest within the same subtype—for example an individual with CLP is more likely to have a sibling with CLP rather than CL or CP— supporting the possibility of subtype-specific effects. Further, “between-subtype” recurrence risks for CL and CLP—for example an individual with CL having a sibling with CLP or vice versa—are lower than within-subtype risks, but are not equal, lending support for the hypothesis that genetic risks for CL and CLP may differ. The lowest recurrence risks were “between-subtype” risks involving CP, but were still higher than the baseline risk in the general population, suggesting some shared etiology between CL/P and CP. In multiplex OFC families with multiple affected individuals, the affected individuals most often all have CL/P or all have CP. Notably, there are also “mixed” families with both CL/P and CP present among relatives, commonly seen with syndromic forms of OFCs.

Taken together these observations imply a genetic predisposition for specific OFC subtypes. While there is limited evidence from genetic association studies supporting subtype-specific risk factors, understanding its granularity has the potential to better inform both biology and diagnosis. The primary focus of the OFC genetics literature has been on CL/P, where over 25 genetic risk loci have been identified to date from genome-wide studies, accounting for a modest portion (~30%) of the overall genetic variance for risk to CL/P (M. J. Dixon et al., 2011; Leslie et al., 2017; Leslie & Marazita, 2013; Ludwig et al., 2017; Yu et al., 2017). By contrast, only one locus has been identified for CP (Leslie, Liu, et al., 2016). To date, subtype-specific associations are limited to three loci: 13q31 near *SPRY2* and *GREM1* (15q13) associated specifically with CLP (Jia et al., 2015; Ludwig et al., 2016; Ludwig et al., 2012), and *GRHL3* (1p36) associated with CP (Leslie, Liu, et al., 2016; Mangold et al., 2016). There is some evidence that markers near *IRF6* (1q32) have a stronger effect on risk for CL than CLP, but this has not been consistently replicated (Leslie & Marazita, 2013; Rahimov et al., 2008).

Given the growing body of evidence suggesting the presence of subtype-specific signals and the broader knowledge base of shared signals, we hypothesize that neither type of statistical signal identifies gene variants that operate in isolation to affect craniofacial development. Rather, it is the combination of shared risk loci and perhaps subtype-specific risk loci affecting an individual’s risk for OFC and their specific OFC subtype. In the current study, we sought to identify novel genetic risk variants for three specific OFC subtypes, CL, CLP, and CP, and examine all genetic risk loci for evidence of being specific to only one OFC subtype or of being shared between two or more subtypes of OFC.

## Results/Discussion

### Genome-wide meta-analysis of CL, CLP, and CP

First, we performed genome-wide meta-analyses for CL, CLP, and CP using imputed genotype data from the GENEVA and Pittsburgh Orofacial Cleft (POFC) consortia (Figure 1, Table S1). The GENEVA consortium used a family-based design and included 461 case-parent trios with CL, 1143 case-parent trios with CLP, and 451 case-parent trios with CP after removing individuals overlapping the two consortia. The POFC consortium included both a case-control arm and a case-parent trio arm, comprising 179 cases and 271 case-parent trios with CL, 644 cases and 1,048 case-parent trios with CLP, 78 cases and 165 case-parent trios with CP, plus 1,700 unaffected controls with no known family history of OFC drawn from the same populations as the unrelated cases. In the POFC case–control subgroup, we used logistic regression to test for association under an additive genetic model and adjusting for 18 principal components of ancestry (Leslie, Carlson, et al., 2016). The two case-parent trio subgroups from POFC and GENEVA were analyzed separately using the allelic transmission disequilibrium test (TDT) (Spielman et al., 1993). The resulting effect estimates for the three analysis groups were combined using an inverse-variance weighted fixed-effects meta-analysis. This procedure was followed separately for genome-wide meta-analyses of CL and CLP (Figure S1); the results of the meta-analysis of CP was previously published (Leslie et al., 2017) and are also depicted in Figure S1.

**Figure 1.**
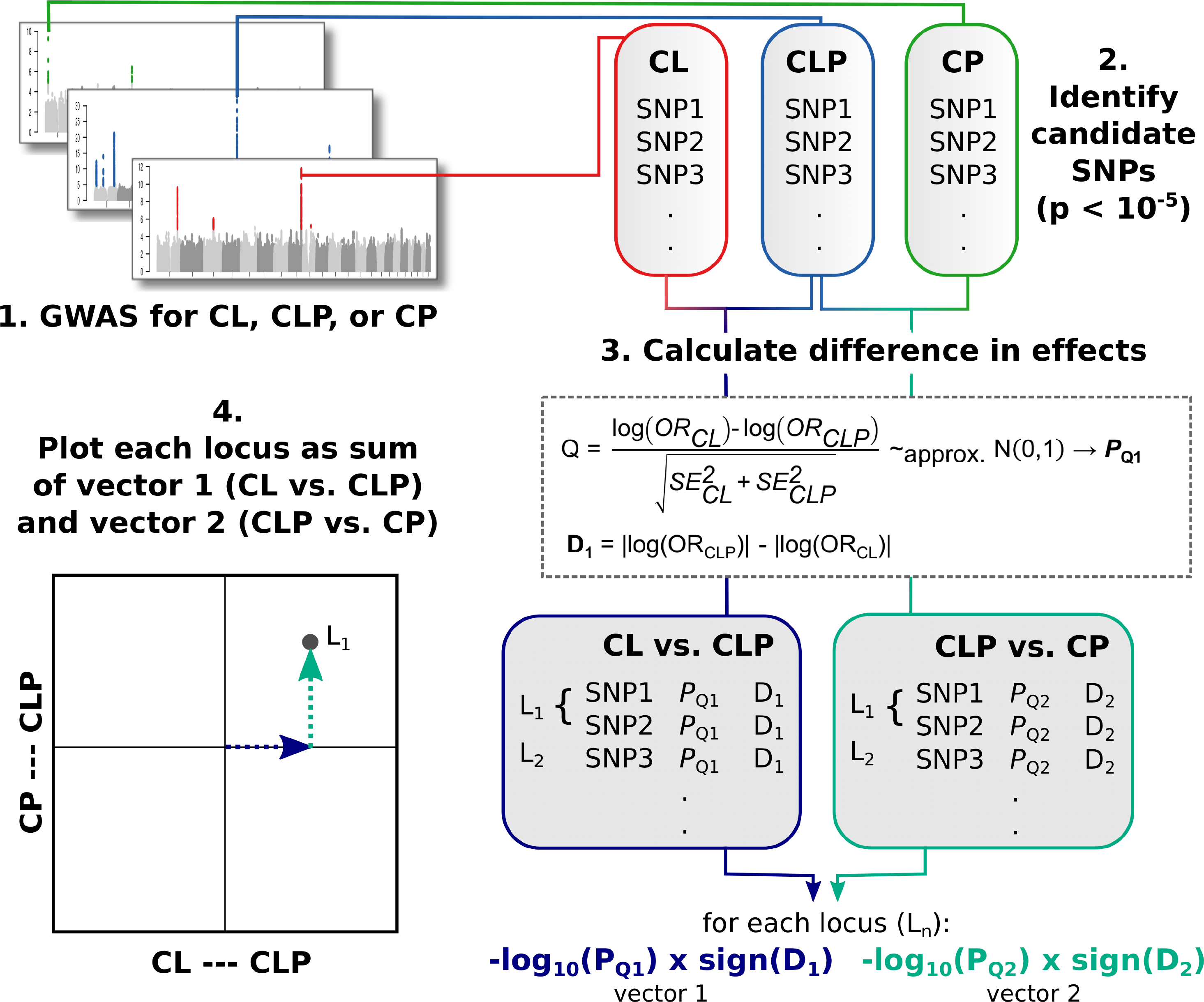
Design and analytical strategy to study phenotypic heterogeneity of orofacial clefts. Analyses consisted of four major steps: (1) GWAS for OFC subtypes, (2) selection of SNPs for analysis (p<10^−5^), (3) calculation of heterogeneity Q-statistic p-values and differences in log odds ratios, and (4) plotting each point as a sum of two vectors, each given by the –log10 p-value of the heterogeneity test times the sign of the direction of effect.

From these three analyses by cleft type, a total of 1,231 SNPs across 29 loci demonstrating suggestive evidence of association (i.e. p<1.00×10^−5^) in any of the three analyses were selected for further follow-up (Figure 1, Table S2). In addition to the 19 recognized risk loci previously reported in GWAS of CL/P or CP separately (Leslie et al., 2017; Yu et al., 2017), this study serves as independent replication for 17q21.3 (near *WNT9B* and *GOSR2*). Nine additional risk loci, although not reaching formal genome-wide significance, were suggested. Three of these—on 1p36.3, 3p14.3, and 5q35.2—have obvious candidate genes previously implicated in craniofacial development or in human craniofacial anomalies. At 1p36.3, a balanced translocation disrupting the *CAPZB* gene was reported in a patient with micrognathia and CP; subsequent studies showed capzb(-/-) zebrafish mutants recapitulated these human phenotypes (Mukherjee et al., 2016). The 5q35.2 signal is adjacent to *MSX2*, a gene critical for human skull development and associated with craniosynostosis, parietal foramina, and orofacial clefting (Wilkie et al., 2000). Finally, the 3p14.3 locus had two independent signals within the same topologically-associated domain containing *WNT5A*, a gene in which mutations can cause mandibular hypoplasia in Robinow syndrome (Hosseini-Farahabadi et al., 2017; Person et al., 2010) and *ERC2*, encoding a synapse protein (Ohtsuka et al., 2002). As *WNT5A* is a stronger candidate gene for OFCs than *ERC2*, we represent the two signals as *WNT5A* “a” and *WNT5A* “b”.

### Identification of independent signals

The 8q24 and *IRF6* loci represent large genomic intervals and have previously shown multiple, statistically independent associations with various OFC phenotypes, although the independence of signals has only been formally tested for the 8q24 gene desert region (Leslie, Carlson, et al., 2016). We separated SNPs into multiple groups based on LD “clumps” calculated with PLINK software (Purcell et al., 2007). In doing so, we confirmed the presence of three independent signals in the *IRF6* region and found evidence for a third signal at the 8q24 region (Figure S2). In total, three loci represented multiple independent signals—1q32 (*IRF6*), 8q24, and 3p14.3 (*WNT5A*)—thus the 29 associated loci comprised 34 total independent signals.

### Comparisons of CL, CLP, and CP

With 1,231 associated SNPs from the 34 independent signals revealed by our three meta-analyses, we used a heterogeneity Q-statistic (Schenker & Gentleman, 2001) to compare effects for each SNP among subgroups to determine if any of these loci showed evidence of subtype-specific or shared risk (Figure 1). Specifically, effects of CL were compared to those of CLP, and the effects of CLP compared to those of CP. These contrasts were selected based on the biological plausibility of shared genetic effects between clefts affecting the lip (CL and CLP) and clefts affecting the palate (CLP and CP). To aid in the identification of subtype-specific variants, the direction of association was calculated by the difference in absolute values of the log odds ratios (i.e. |log(OR_CLP_)| - |log(OR_CL_)|, |log(OR_CLP_)| - |log(OR_CP_)|).

In our application of this approach, some components of the Q-statistic (i.e. the odds ratio for each cleft subtype) are inherently correlated because we have a shared pool of controls in a subset of the study. This dependence among the effect estimates is potentially problematic because the variants that were selected for comparison between CL, CLP, and CP subtypes, were selected because of their marginal association with at least one subtype. To account for this, we conducted a permutation procedure for the difference in log odds ratios for each comparison (i.e. the numerator of the Q-statistic), wherein we randomized the cleft type definitions among cases (and trio probands) and performed the association testing procedure with the permuted cleft subtypes. We then generated an empirical distribution of the difference in log odds ratios between CLP and CL and between CLP and CP from up to 10,000 permutations, and used this to calculate an empirical p-value. The results of the permutation procedure are given in Table S3. The resulting empirical p-values were largely similar to the original p-values that did not account for the shared controls (Figure S3),so the results, figures and tables that follow are henceforth based onfrom these original Q-statistic p-values.

We sought to visually represent these findings so the evidence of subtype-specific effects become clear. Rather than dichotomizing genetic effects as “subtype-specific” or “shared”, we wanted to represent both the statistical evidence for heterogeneity and the overall statistical evidence of association with cleft subtypes for each locus. To this end, we developed a graphical representation, hereafter referred to as “the cleft map” to describe the statistical effect of numerous genetic loci on the architecture of OFCs. On the cleft map, the position of any single SNP is determined by the sum of two vectors, each given by the –log_10_ p-value of the heterogeneity Q-statistic times the sign of the direction of the locus (Figure 1). The position of a SNP thus represents how heterogeneous its estimated effects were between two cleft subtypes; SNPs further from the origin demonstrate more statistical evidence of heterogeneity. The x-axis of the cleft map represents the CL vs. CLP comparison and the y-axis represents the CLP vs. CP comparison.

When all SNPs for a locus were plotted, they generally clustered in specific locations based on the cleft type(s) with which the locus is associated (Figure 2, Figure S4). For example, SNPs located near the origin of the axes were those that showed less evidence of cleft-subtype specificity. Such a pattern occurred for all 4 tested SNPs at the *FOXE1* locus (Figure 2A), consistent with existing literature indicating associations with all OFC subtypes (Leslie et al., 2017; Moreno et al., 2009). In contrast, all 13 SNPs at the *GRHL3* locus (Figure 2B) showed a significant CP-specific association (p_CLP.CP_<6.2×10^−5^) and are positioned along the y-axis in the lower half of the map. Similarly, all 126 SNPs in the *IRF6* “a” locus (Figure 2C) showed a significant association with CL/P, as evidenced by significant differences in the CLP-CP comparison (p_CLP.CP_<6.9×10^−5^), but there was no evidence of difference between CL and CLP (p_CLP.CL_>0.26). The independent *IRF6* “a” and *IRF6* “b” regions contain rs2235371 (Zucchero et al., 2004) and rs642961, respectively. Previous studies suggested that rs642961 was preferentially associated with CL (Rahimov et al., 2008). Although *IRF6* “b” SNPs show a quite complex relationship with different cleft types probably related to underlying differences in linkage disequilibrium (LD) (Figure S2), the location of the SNP cluster containing rs642961 in the upper-left quadrant of the cleft map supports a stronger effect on risk for CL compared to CLP (Figure 2D).

**Figure 2.**
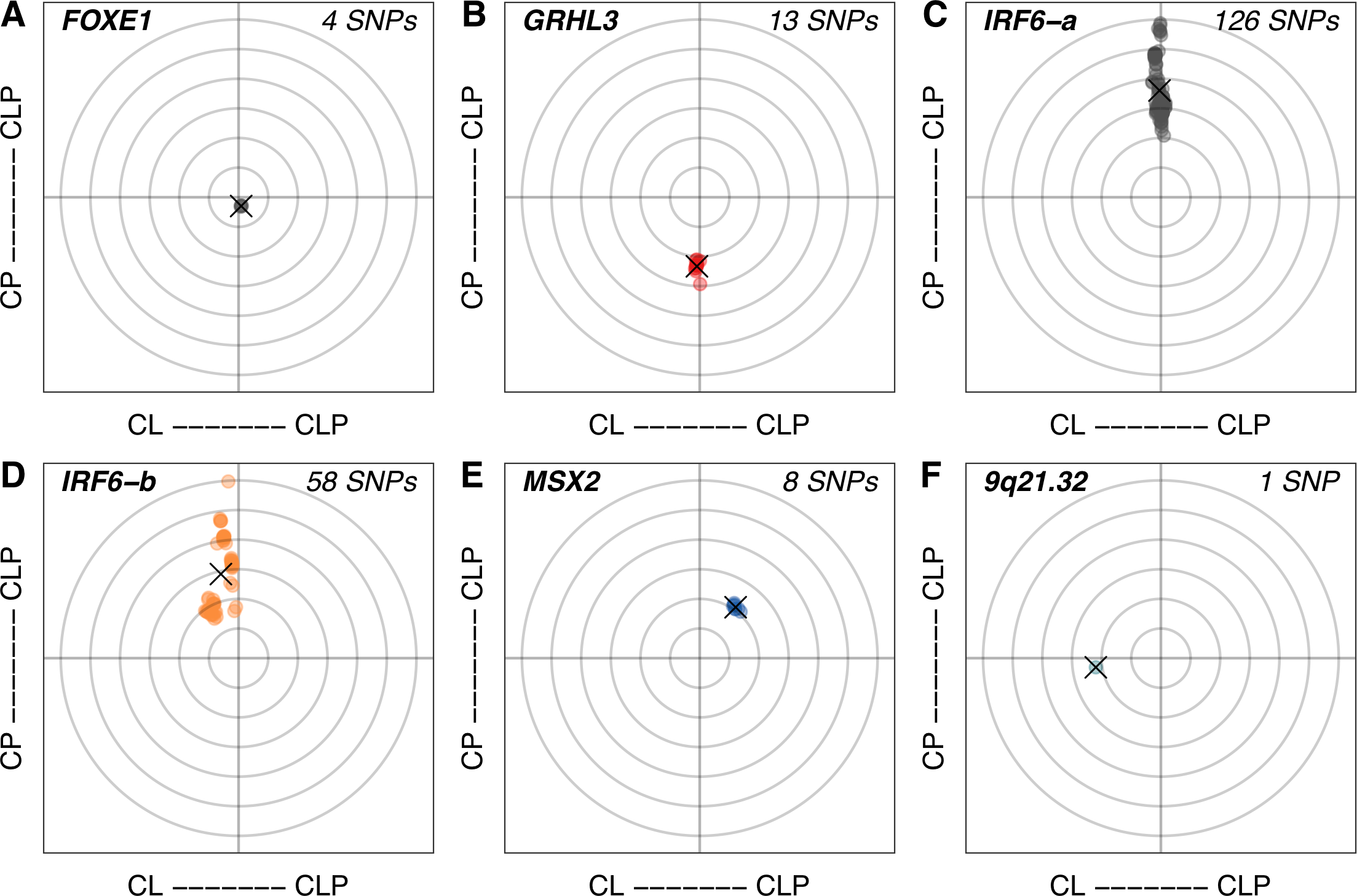
Subtype effects for SNPs at representative loci. For each SNP per locus, the effects for CL and CLP, and CLP and CP were compared with heterogeneity Q-statistics. The direction of association was determined by the difference in absolute values of the log odds ratios (i.e. |log(OR_CLP_)| - |log(OR_CL_)|, |log(OR_CLP_)| - |log(OR_CP_)|). The coordinates of each SNP were determined by the sum of two vectors, each given by the –log10 p-value of the Q statistic times the sign of the direction. The x-axis of the cleft map represents the CL vs. CLP comparison and the y-axis represents that CLP vs. CP comparison. Concentric circles around the origin based on p-values of the Q-statistics are given for reference (0.01, 0.0001, increasing by 10^−2^). The centroid of each cluster of SNPs is represented by an “X”.

With a few exceptions, the tight clustering of SNPs allowed us to simplify the map by representing each locus region centered around its SNP cluster (Table 1, Figure 3). In addition to the loci described above, several others demonstrated some evidence of cleft subtype specificity including *MSX2* with CLP (Figure 2E, p_CLP.CP_<7.3×10^−4^, p_CLP.CL_<5.6×10^−3^), *WNT5A “a”* with CLP (Figure S4F, p_CLP.CP_<0.09, p_CLP.CL_<1.1×10^−4^), 9q21.32 with CL (Figure 2F, p_CLP.CP_=0.24, p_CLP.CL_=4.06×10^−5^), and 5p13.2 with CP (Figure S4K, p_CLP.CP_<2.2×10^−4^).

**Table 1.** Summary of Cleft Map results for all 34 independent signals across 29 loci

| Region | Candidate Gene(s) | CL vs. CLP |  | CLP vs. CP |  | Centroid Coordinates | N SNPs |
| --- | --- | --- | --- | --- | --- | --- | --- |
|  |  | Minimum P-value from Q statistic | Maximum P-value from Q statistic | Minimum P-value from Q statistic | Maximum P-value from Q statistic |  |  |
| 1p36.13 | PAX7 | 0.015 | 0.785 | **7.65E-09** | 0.014 | 0.9917,3.326 | 73 |
| 1p36.13 | CAPZB | 0.034 | 0.133 | 9.38E-04 | 5.00E-03 | 1.188,2.679 | 15 |
| 1p36.11 | GRHL3 | 0.483 | 0.969 | **1.36E-06** | **6.24E-05** | -0.191,-4.663 | 13 |
| 1p22 | ARHGAP29 | 0.222 | 0.975 | **6.26E-04** | 0.159 | 0.1046,1.473 | 16 |
| 1q23.1 | ETV3 | 0.010 | 0.014 | 0.057 | 0.068 | 1.914,1.214 | 6 |
| 1q32 "a" | IRF6 | 0.262 | 0.979 | **1.78E-12** | **6.93E-05** | -0.0933,7.207 | 126 |
| 1q32 "b" | IRF6 | 5.46E-03 | 0.672 | **1.18E-12** | 2.16E-03 | -1.199,5.751 | 58 |
| 1q32 "c" | IRF6 | 9.44E-02 | 0.994 | **7.17E-09** | 3.43E-02 | 0.1599,1.256 | 126 |
| 2p24.2 | FAM49A | 0.510 | 0.991 | 0.092 | 0.196 | 0.1883,0.8341 | 53 |
| 3p14.3 "a" | WNT5A | **8.19E-05** | **1.16E-04** | 0.060 | 0.090 | 4.011,1.134 | 2 |
| 3p14.3 "b" | WNT5A | 0.057 | 0.078 | 0.207 | 0.273 | 1.194,0.632 | 5 |
| 3q12.1 | COL8A1,<br>FILIP1L | 2.72E-03 | 0.014 | 1.60E-03 | 0.054 | -2.308,1.964 | 106 |
| 3q28 | TP63 | 0.580 | 0.933 | 8.75E-03 | 0.018 | -0.1078,1.871 | 5 |
| 4q21.1 | SHROOM3 | 0.708 | 0.995 | **5.20E-05** | 2.95E-03 | 0.02269,3.2 | 13 |
| 5p13.2 | CAPSL,<br>SKP2 | 0.736 | 0.933 | **2.00E-05** | **2.29E-04** | -0.03216,-4.047 | 4 |
| 5q35.2 | MSX2 | 1.75E-03 | 0.006 | **1.92E-04** | **7.36E-04** | 2.421,3.437 | 8 |
| 6p22 | TRIM10 | 0.332 | 0.332 | 0.361 | 0.361 | 0.4784,0.4425 | 1 |
| 8q21 | DCAF4L2,<br>MMP16 | 0.305 | 0.916 | 1.59E-03 | 0.254 | 0.1946,1.609 | 10 |
| 8q22.3 | FZD6,<br>RIMS2 | 0.012 | 0.012 | 0.136 | 0.136 | 1.931,0.8674 | 1 |
| 8q24 "a" |  | 0.237 | 0.974 | **2.10E-09** | 0.003 | -0.2666,5.396 | 57 |
| 8q24 "b" |  | 2.87E-03 | 0.992 | **3.72E-07** | 0.088 | -0.7306,3.179 | 108 |
| 8q24 "c" |  | 0.092 | 0.980 | **7.02E-05** | 0.024 | 0.1114,2.671 | 94 |
| 9q21.32 | TLE1 | **4.06E-05** | **4.06E-05** | 0.240 | 0.240 | -4.391,-0.6196 | 1 |
| 9q22 | FOXE1 | 0.602 | 0.708 | 0.227 | 0.271 | 0.174,-0.593 | 4 |
| 10q25 | VAX1,<br>SHTN1 | 0.316 | 0.921 | 0.179 | 0.415 | 0.2362,0.5786 | 22 |
| 12q13 | KRT8,<br>KRT18 | 0.088 | 0.573 | **1.10E-05** | 1.50E-03 | 0.5406,3.777 | 12 |
| 13q31 | SPRY2 | 0.096 | 0.447 | **9.08E-05** | 1.36E-03 | 0.7439,3.599 | 10 |
| 13q32.3 | CLYBL,<br>ZIC5, ZIC2 | 0.109 | 0.271 | 0.062 | 0.586 | 0.7145,0.8081 | 13 |
| 15q22.2 | TPM1 | 0.261 | 0.261 | 0.059 | 0.059 | 0.5834,1.231 | 1 |
| 15q24.1 | ARID3B | 0.519 | 1.000 | 3.61E-03 | 0.015 | 0.08757,2.12 | 137 |
| 17p13 | NTN1 | 0.020 | 0.953 | **1.28E-06** | 0.391 | 0.6106,2.531 | 42 |
| 17q21.32 | WNT9B | 0.357 | 0.707 | 0.239 | 0.704 | 0.3152,0.3496 | 39 |
| 17q22 | NOG | 0.460 | 0.475 | **1.26E-06** | **7.56E-06** | -0.3303,-0.3891 | 2 |
| 20q12 | MAFB | 0.071 | 1.000 | 2.13E-03 | 0.389 | 0.6294,1.634 | 42 |

**Figure 3.**
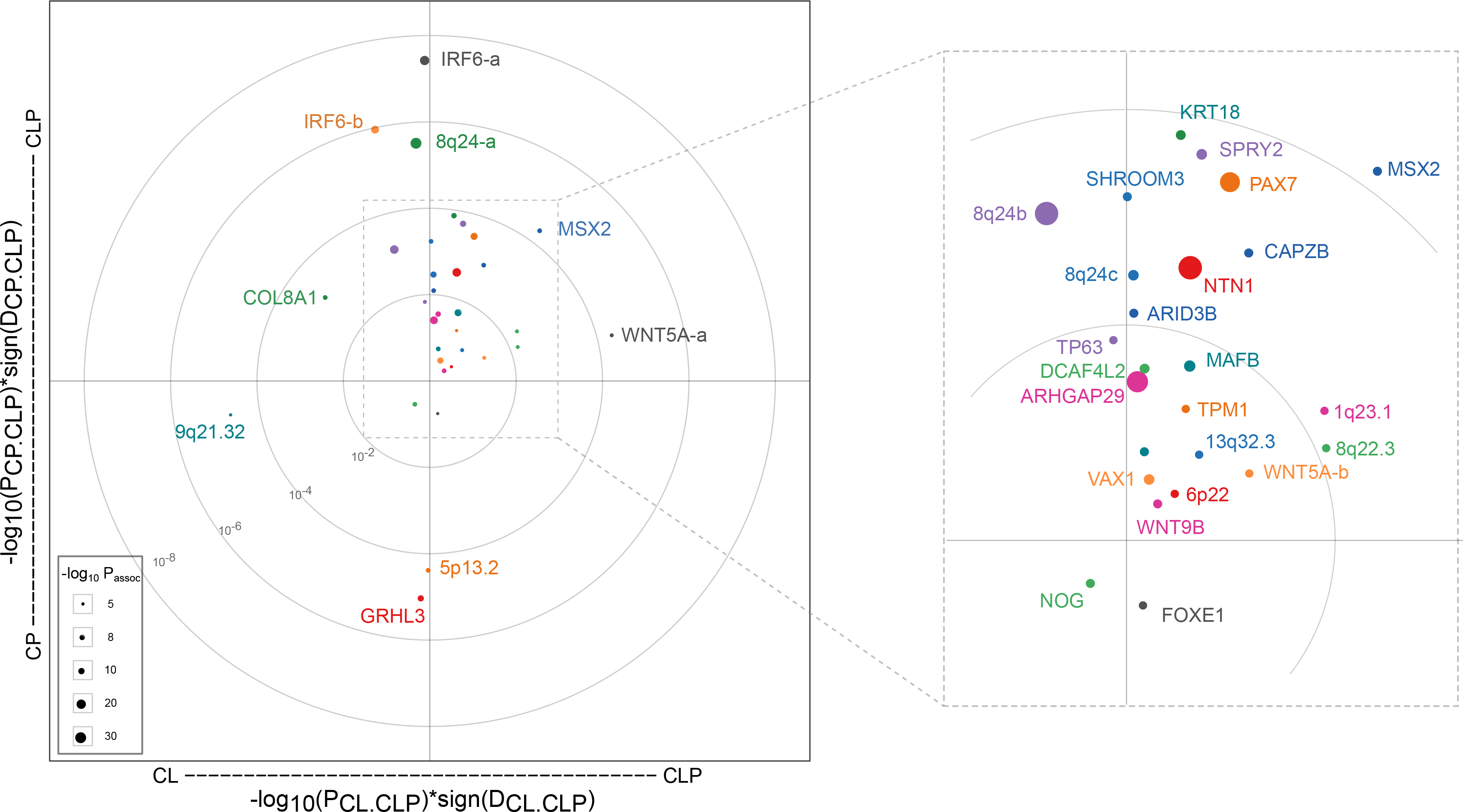
The Cleft Map. Each of the 29 loci are represented by a single point as the centroid of all SNPs at the locus. The size of the point is scaled to the –log10 p-value for the most significant SNP in the meta-analyses of CL, CLP, and CP. Concentric circles about the origin based on p-values of the Q-statistics are given for reference (0.01, 0.0001, increasing by 10^−2^). Point size is scaled to represent the best p-value observed in the meta-analyses. Points are colored for clarity of gene name labels.

### Expression analyses in associated loci

Above we have referred to each associated locus by either a plausible candidate gene (e.g., *IRF6*) based on the literature on OFCs and craniofacial development or as a genetic location (e.g., 8q24) for gene deserts or new loci. Although this comports with the standards of the field, recent work demonstrates that disease-associated variants can regulate distant genes, suggesting that the nearby genes prioritized by GWAS may not always be involved (Gupta et al.; Smemo et al.). In addition, multiple genes in a given region may be co-regulated and expressed in similar tissues or expressed in distinct compartments within the craniofacial complex (Attanasio et al., 2013). We, therefore, wanted to agnostically explore gene expression profiles of all genes contained in these regions to determine if their expression profiles could explain clustering of associated genes or provide mechanistic insights into the pathogenesis of OFC subtypes.

We first identified all of the genes located in the same topologically associated domains (TADs), because the associated SNPs may have regulatory functions and there is evidence that such SNPs are more likely to act upon genes located within their own TAD (Table S4) (Dekker et al., 2013). We used published topological data from human embryonic stem cells because although there are no known craniofacial-specific TADs, boundaries are largely conserved across cell types (J. R. Dixon et al.). To prioritize highly expressed genes, we identified those genes with mouse homologs reported to be differentially expressed among several transcriptomic datasets from key periods, regions, and tissues in mouse facial development (Hooper et al., 2017). Hooper et al. previously integrated these datasets with a weighted gene co-expression network analysis to generate 75 co-expression modules describing gene expression in the developing mouse face. We used these modules to annotate each gene in our list (Table S5). Of the 222 homologs in our 29 loci, 101 (45.44%) were present among these co-expression modules. Each region contained at least one gene assigned to a co-expression module, but only seven regions contained a single gene or only one gene with documented craniofacial expression (Figure S6). Overall, this is not surprising given these co-expression modules contained over 8,000 genes, or approximately 30% of the mouse genome.

To further refine the possible candidate genes, we turned to a complementary resource of gene expression profiles, SysFACE (<u>Sys</u>tems tool for cranio<u>fac</u>ial <u>e</u>xpression-based gene discovery) (Table S6) (Liu et al., 2017), which allows easy visualization of data from orofacial tissue microarrays or RNA-seq datasets for mandible, maxilla, frontonasal prominence, and palate, collected largely as part of the FaceBase consortium (Brinkley et al., 2016; Hochheiser et al., 2011). We used SysFACE to examine enriched craniofacial expression by comparing orofacial tissue data with embryonic whole body tissue (Lachke et al., 2012). For most loci, the SysFace analysis corroborated the co-expression modules or prioritized one gene. For example, at the *PAX7* locus, although three genes (*Pax7, Klhdc7a*, and *Aldh4a1*) were found in craniofacial co-expression modules only *Pax7* showed strong enrichment in SysFACE.

For the six new loci without any clear candidate genes (5p13.2, 6p22, 8q22.3, 9q21.32, 13q32.3), we used both datasets to identify likely candidate genes. At the 5p13.2 locus, *Skp2* was present in the mesenchyme expression module and showed high SysFACE scores across multiple processes in the microarray data and RNA-seq expression in the palate. Other genes (*Capsl* and *Slc1a3*) were in the ectoderm module with enriched expression in the palate. Because this locus is associated only with CP, any or all of these genes could be relevant. At the 6p22 and 8q22.3 loci, multiple genes were found in both the Hooper and SysFACE datasets, but were expressed in different expression modules. Although *FZD6* (8q22.3) was previously implicated in CLP by linkage in a large multiplex CLP family, as well as by craniofacial anomalies observed in a *fzd6* morphant (Cvjetkovic et al., 2015), it is possible multiple genes contribute to the association signals at these loci. In sum, each newly associated locus contained one or more genes with craniofacial expression; detailed *in vivo* analyses will be required to pinpoint specific causal genes.

We performed an enrichment analysis for the set of genes present in these co-expression modules. These genes were enriched for broad biological processes such as “embryonic morphogenesis”, “epithelium development” and human and mouse phenotypes related to OFCs, including “oral cleft”, “perinatal lethality”, “abnormal craniofacial morphology”. Unfortunately, the broad terms from the enrichment analysis did not support more specific hypotheses about the pathogenesis of OFCs. The co-expression modules revealed a critical role for ectodermal genes in OFC pathogenesis, and fewer loci with mesenchymally-expressed genes. Most genes, however, were broadly expressed across multiple facial prominences, limiting our ability to hypothesize about any one mechanism for how these genes relate to OFC subtypes, but these tools will be useful for prioritizing genes for future association studies.

A few genes, however, had very specific expression patterns worthy of further discussion. As one example, *PAX7* expression was restricted to the frontonasal prominence, whereas other loci (i.e., *SPRY2*, *MSX2*) that clustered nearby in the cleft map showing stronger evidence of effects on risk for CLP were more broadly expressed in the maxilla and mandible. Similarly, the *COL8A1* locus showed a stronger effect on CL than CLP, but was still very strongly expressed in the palate. Interestingly, *Col8a1* expression was enriched early at E10.5 in the frontonasal and maxillary prominences, when lip fusion takes place. These patterns are consistent with the direction of the SNP effects at this locus where the same alleles conferred a protective effect (OR < 1) for CL and CLP scans, but a modest, (and not formally significant) risk effect (OR > 1) in the CP scan. Thus, our data argue both the early expression and the palatal expression are important. It is possible that the as-yet-unknown functional SNPs could promote palatal fusion and protect against a CP; alternatively, they could dysregulate *COL8A1* promoting ectopic expression in the lip. Our current study cannot definitively answer these questions, but by demonstrating this locus may exert a stronger effect on risk to CL while the gene is expressed in the palate, it could motivate more targeted follow-up studies.

In our statistical analyses, we observed most of the loci previously identified in CL/P GWAS were positioned along the y-axis, indicating that there was no statistical difference between CL and CLP. This is not surprising, as a combined study of CL and CLP has the greatest power to identify loci with similar effects in both of these subgroups. One limitation of this study is that we did not capture several loci discovered in previous CL/P GWAS (e.g., *RHPN2* (Leslie, Carlson, et al., 2016)) because no SNP showed p-values better than 10^−5^. As these loci were found only in the combined CL/P group, we would expect them to be positioned along the y-axis. Such SNPs show the statistical power of traditional analyses of CL/P, which have been successful. However, our study also demonstrates there are multiple loci with subtype-specific effects (e.g. *WNT5A*) or with stronger effects for CL than CLP (e.g. *COL8A1*) that are more difficult to detect in the combined analyses.

An important contribution of this work was the careful examination of the three large loci with multiple, independent signals based on LD. Of these, we found evidence for independent signals within *WNT5A* and *IRF6* exerting potentially different effects between subtypes. In contrast, the three 8q24 signals were largely overlapping and associated with the combined phenotype CL/P (Figure S4 M-O); only the 8q24 “c” signal showed even marginal evidence of a stronger effect with CL than CLP. At the *IRF6* locus, the “b” signal was tagged by rs642961, a SNP that disrupts the binding site of TFAP2α in the MCS9.7 enhancer (Rahimov et al., 2008). However, MCS9.7 activity did not completely recapitulate endogenous *IRF6* activity, most notably in the medial-edge epithelium during palatal fusion, indicating the presence of some other enhancer (Fakhouri et al., 2012). Similarly, the 8q24 gene desert contains multiple craniofacial enhancers (Attanasio et al., 2013) which influence *Myc* expression (Uslu et al., 2014). Enhancers are known to have restricted activity patterns and often act in a modular fashion to control gene expression by activating expression in different anatomical regions or at different points during development (Visel et al., 2009). These characteristics present a logical mechanism to drive phenotypic heterogeneity. Understanding the logic of these enhancers, the function of SNPs within them, and the relationship between risk alleles and disease subtypes will be critical for fully understanding the etiology of OFCs.

Our approach works well to describe SNPs and loci where the direction of the effect is the same among subtypes. However, when the odds ratios exhibit different effect directions, the log odds ratios may be close in value, and the sign representing the direction of difference in effects may fluctuate according to the subtype with a slightly larger effect at any given SNP. It is possible to diagnose these instances by plotting estimated effects of the individual SNPs which resulting in two clusters of SNPs positioned on opposite sides of the plot (as seen in Figure S4Z). Overall, plotting the centroid (as shown for the *NOG* locus (Figure S4AA)) most closely represents the overall picture of the locus—i.e. that there is no association with one particular cleft type. However, if an allele truly increases risk for one cleft type and reduces risk for another, as was recently reported for *NOG* (Moreno Uribe et al., 2017), this visualization may be obscuring biologically meaningful results. Similarly, we note the presence of a set of SNPs within *IRF6* “c” (apparently independent of the *IRF6* “a” and *IRF6* “b” signals), with p-values in the CP meta-analysis of ~8×10^−3^ and whose minor alleles appear to increase risk for CP; these same alleles appear to decrease risk for CL/P. Although genetic association studies overwhelmingly support an association between common SNPs in *IRF6* and risk of CL/P, dominant mutations in the gene cause Van der Woude syndrome, recognized as one of few syndromes where both CL/P and CP occur within the same family. With this new information about this locus, it may be time to revisit the idea that common SNPs within this locus could act as modifiers of which OFC phenotype appears in Van der Woude families (Leslie et al., 2013).

## Conclusions

In summary, we developed an approach to dissect and visualize the genetic contribution to phenotypic heterogeneity of OFCs. Overall, our results point to shared effects for most GWAS loci between CL and CLP. However, we identified novel genetic associations, showing evidence of subtype specificity that would have been missed by traditional analyses where CL and CLP or all three OFC subtypes are combined. In addition, we were able to identify and analyze three loci (*IRF6*, *WNT5A*, and 8q24) that contain multiple independent signals. Ours is the first study to formally confirm multiple signals from *IRF6* locus, which had been suggested previously (Rahimov et al., 2008). Importantly, some of these signals showed different effects on the different OFC subtypes, adding an additional layer of complexity to the genetic architecture of this most common group of craniofacial malformations. Finally, cross-referencing our results with gene expression data has generated new hypotheses about mechanisms by which OFC subtypes may occur.

In this work, we focused only on fetal contributors to OFC risk, but recognize that genetic studies alone are unlikely to completely elucidate the etiology of OFCs or other complex traits where the etiology likely includes significant environmental components, epigenetic factors, parent-of-origin effects, stochastic processes, or additional genetic modifiers. With respect to OFCs, we have previously shown evidence of both common and rare genetic modifiers possibly distinguishing between CL from CLP (Carlson, Standley, et al., 2017; Carlson, Taub, et al., 2017). Our approach may serve to guide targeted tests for gene-gene interaction and risk score analyses to further disentangle the complex etiologic architecture of OFCs. Future extensions of this approach can incorporate these modifiers and interactions, and can examine other subtype definitions, other structural birth defects, or any other complex trait with phenotypic heterogeneity where subtypes can be delineated. Finally, we note our approach relies on GWAS or candidate gene study summary statistics. In our cleft map application, the summary statistics were derived from analyses using shared controls so a permutation procedure using individual-level data was used to verify results. However, when effect estimates can be derived from independent associations, this procedure can use summary statistics alone to easily leverage information from multiple studies to dissect the complex architecture of heterogeneous traits.

## Methods

### Contributing GWAS studies

Two consortia contributed to this study. The first, from the GENEVA consortium, used a family-based design and included 461 case-parent trios with cleft lip (CL), 1,143 case-parent trios with cleft lip and palate (CLP), and 451 case-parent trios with CP, respectively, from populations in Europe (Denmark and Norway), the United States, and Asia (Singapore, Taiwan, Philippines, Korea, and China). The specifics of this study have been described previously (Beaty et al., 2010; Leslie, Carlson, et al., 2016; Leslie, Liu, et al., 2016). Briefly, samples were genotyped for 589,945 SNPs on the Illumina Human610-Quadv.1_B BeadChip, genetic data were phased using SHAPEIT, and imputation was performed with IMPUTE2 software to the 1000 Genomes Phase 1 release (June 2011) reference panel. Genotype probabilities were converted to most-likely genotype calls with the GTOOL software (http://www.well.ox.ac.uk/~cfreeman/software/gwas/gtool.html).

The second consortium included samples from the Pittsburgh Orofacial Cleft (POFC) study, comprising 179 cases and 271 case-parent trios with CL, 644 cases and 1,048 trios with CLP, 78 cases and 165 trios with CP, and 1,700 unaffected controls with no history of craniofacial anomalies. Participants were recruited from 13 countries in North America (United States), Central or South America (Guatemala, Argentina, Colombia, Puerto Rico), Asia (China, Philippines), Europe (Denmark, Turkey, Spain), and Africa (Ethiopia, Nigeria). Additional details on recruitment, genotyping, and quality controls were previously described (Leslie, Carlson, et al., 2016; Leslie, Liu, et al., 2016). Briefly, these samples were genotyped for 539,473 SNPs on the Illumina HumanCore+Exome array. Data were phased with SHAPEIT2 and imputed using IMPUTE2 to the 1000 Genomes Phase 3 release (September 2014) reference panel. The most-likely genotypes (i.e. genotypes with the highest probability [Q]) were selected for statistical analysis only if the genotype with the highest probability was greater than 0.9. A total of 412 individuals were in both the GENEVA OFC and POFC studies, so we excluded these participants from the GENEVA study for this analysis. Informed consent was obtained for all participants and all sites had both local IRB approval and approval at the University of Pittsburgh, the University of Iowa, or Johns Hopkins University. Individual level genotype and phenotype data for the GENEVA and POFC studies are available from dbGaP: phs000774.v1.p1 and phs000094.v1.p1.

### Genome-wide meta-analyses for CL, CLP, and CP

We previously described our methods for quality control and meta-analysis of the GENEVA OFC and POFC studies (Leslie et al., 2017) and quality control procedures were completed in each contributing study and were described extensively in the original publications (Leslie et al. (2016b), Leslie et al. (2016a), Beaty et al. (2010) and Beaty et al. (2011)). Briefly, SNPs with minor allele frequencies (MAF) less than 1% or those deviating from Hardy-Weinberg Equilibrium (HWE p < 0.0001) in the parents or control subject were excluded from the analysis. To account for different marker sets and identifiers between the two imputed datasets, the final analysis included only those overlapping SNPs that were matched on chromosome, nucleotide position, and alleles. GWAS was performed for CL, CLP, and CP separately on SNPs with minor allele frequencies (MAF) greater than 1% and not deviating from Hardy-Weinberg Equilibrium (HWE p > 0.0001) in the parents or control subjects. Tests of association using unrelated cases and controls matched for population of origin used logistic regression models under an additive genetic model while including 18 principal components of ancestry (generated via principal component analysis of 67,000 SNPs in low linkage disequilibrium across all ancestry groups) to adjust for population structure. Case-parent trios were analyzed with the allelic transmission disequilibrium test (TDT) implemented in PLINK. Within each OFC subtype, the resulting effects estimates were combined in an inverse-variance weighted fixed-effects meta-analysis. The combined estimate, a weighted log odds ratio, should follow a chi-squared distribution with two degrees of freedom under the null hypothesis of no association.

### Comparisons of cleft types

From the three meta-analyses, SNPs demonstrating suggestive evidence of association (i.e. p < 1.00×10^−5^) in any scan were considered for further analysis. For each SNP, two heterogeneity Q-statistics were calculated to compare effects of CL to CLP, and CLP to CP (Schenker & Gentleman, 2001). The magnitude of the log odds ratios for these two cleft subtypes were also compared to indicate the direction of effect (i.e. which subtype showed a stronger effect). Together, the Q-statistics and directions of effect for the two comparisons (CL to CLP, CLP to CP) prescribe a point for each SNP on the cleft map. For each region, the centroid of these points was calculated to inform the overall effect of the locus.

There is an inherent dependence between odds ratios for each cleft subtype that arises due to shared controls in each of the three case-control components of the meta-analyses. Because of this, we performed a sensitivity analysis on the Q-statistic comparison using a permutation procedure. For each permutation, we randomly shuffled the cleft subtypes among the cleft cases and re-performed the meta-analysis for CL, CLP, and CP separately, and calculated the difference in log odds ratios for each comparison (CLP with CL and CLP with CP). We used the permuted results to generate an empirical null distribution of the difference in log odds ratios and to calculate an empirical p-value. This permutation procedure was computationally expensive, so large scale permutations were infeasible. So, we performed the permutations in an adaptive fashion. For each variant, a minimum of 100 permutations were performed. Then a naïve confidence interval for the true p-value was calculated using the resulting permutations. If that confidence interval contained zero, the maximum number of permutations was increased 10-fold. This was conducted iteratively until the naïve confidence interval did not contain zero, or the maximum number of permutations (10,000) was reached. We note that the smallest empirical p-value is then 0.0001 (i.e. 1 in 10,000), limiting the usefulness of this permutation procedure for loci with very significant differences in effects across cleft subtypes.

### SysFACE: mouse orofacial transcriptome data analysis

Mouse orthologs of human candidate genes were analyzed for their absolute expression and enriched expression in orofacial tissue microarray datasets on the Affymetrix 430 2.0 platform (GeneChip Mouse Genome 430 2.0 Array) deposited in NCBI GEO (https://www.ncbi.nlm.nih.gov/geo/) and FaceBase (https://www.facebase.org) (Table S6). Enriched expression was estimated by comparing orofacial tissue data with whole embryonic body tissue (WB) reference control obtained from series (GSE32334) as previously described (Lachke et al., 2012). Datasets for mandible (mouse embryonic stages E10.0, E10.5, E11.0, E11.5, E12.0, E12.5), maxilla (E10.5, E11.0, E11.5, E12.0, E12.5,) and frontonasal (E10.5, E11.0, E11.5, E12.0, E12.5) were obtained from the series GSE7759. Palate datasets were obtained for mouse E13.5 (FaceBase series: FB00000468.01), E14.5 (FB00000474.01, GSE11400) and P0 (GSE31004). RNA-seq data on Illumina HiSeq2500 platform for mouse E14.5 posterior oral palate (FB00000768.01), anterior oral palate (FB00000769.01) and WB (whole body, unpublished) were also used. ‘R’ statistical environment (http://www.r-project.org/) was used to import raw microarray datafiles on Affymetrix 430 2.0 platform, followed by background correction and normalization using Affy package (Gautier et al., 2004) available at Bioconductor (www.bioconductor.org). Using a AffyBatch function, present/absent calls for probe sets were calculated and those with the highest median expression at significant p-values were collapsed into genes (Gautier et al., 2004). Differential gene expression (DEGs) and enrichment scores for all four orofacial tissues compared to WB were calculated using limma (Ritchie et al., 2015), and the detailed microarray workflow is described elsewhere (Anand et al., 2015). RNA-seq data on Illumina HiSeq2500 platform were first subjected to quality control analysis for reads by using FastQC (http://www.bioinformatics.babraham.ac.uk/projects/fastqc/), and then subjected to sequence trimming and clipping using Trimmomatic (Bolger et al., 2014) with in house scripts (https://github.com/atulkakrana/preprocess.seq) (Mathioni et al., 2016). Reads were aligned against the *Mus musculus* reference genome using TopHat v2.0.9 (Trapnell et al., 2009) with recommended settings. Transcript assembly for measuring relative abundances was performed using Cufflinks v2.1.1 (Trapnell et al., 2009). After merging the assemblies by the function Cuffmerge, DEGs were identified using the Cuffdiff function (Trapnell et al., 2009). Statistically significant DEGs (comparison of orofacial datasest with WB) were identified using an in-house Python script.

## Grant Numbers

This work was funded by grants from the National Institutes of Health: R00-DE025060 [EJL], R03-DE024776 [SAL, IS], X01-HG007485 [MLM, EF], R01-DE016148 [MLM., SMW], R01-DE009886 [MLM], R21-DE016930 [MLM], R01-DE014667 [LMM], R01-DE012472 [MLM], R01-DE011931 [JTH], R01-DE011948 [KC], R37-DE008559 [JCM, MLM], U01-DD000295 [GLW], R00-DE024571 [CJB], S21-MD001830 [CJB], U54-MD007587 [CJB], R00-DE022378 [AB], R01-DE014581 [TB], U01-DE018993 [TB], and Robert Wood Johnson Foundation Grant #72429 [AB]. Genotyping and data cleaning were provided via an NIH contract to the Johns Hopkins Center for Inherited Disease Research: HHSN268201200008I.

## Acknowledgments

We thank the study participants for their enthusiastic participation and acknowledge the local recruitment staff and collaborators for their tireless efforts that made this study possible.

## Supporting Information Captions

**Figure S1. Results of genome-wide meta-analyses for (A) cleft lip (CL), (B) cleft lip and palate (CLP), and (C) cleft palate (CP).** SNPs with p-values less than 1.00×10^−5^ are highlighted.

**Figure S2. Identifying independent signals with LD clumping.** Results from our previously published CL/P meta-analysis were analyzed with the PLINK (Purcell et al., 2007) clumping procedure (--clump) The clumping procedure takes all SNPs that are significant at the threshold of the index SNPs and forms clumps of all other SNPs that are within 250kb from the index SNP and that are in linkage disequilibrium with the index SNP, based on an r-squared threshold of 0.5. The PLINK clump command is a greedy algorithm so each SNP will only appear in a single clump. To simplify the clumps, we combined clumps. (A) The IRF6 “a” signal consists of only SNPs in “clump 1”; the IRF6 “b” signal consists of SNPs from clumps 2-4; the IRF6 “c” signal consists of SNPs from clumps 5 and 6. (B) The 8q24 “a” signal consists of SNPs in clumps 1-3; the 8q24 “b” signal consists of SNPs from clumps 4 and 5; 8q24 “c” consists of SNPs from clumps 6 and 7. Panels (C) and (D) show the cleft map plots separately for each clump for IRF6 and 8q24, respectively. Concentric circles indicate significance thresholds for 0.01, 0.001, 0.0001, 1×10×^−6^, 1×101×10×10^−8^, 1×10^−10^, and 1×10^−12^.

**Figure S3. Comparison of p-values obtained from the original Q statistic and empirical p-values derived from 10,000 permutations**.

**Figure S4. Cleft map for each locus.** Concentric circles indicate significance thresholds for 0.01, 0.001, 0.0001, 1×10×^−6^, 1×101×10×10^−8^, 1×10^−10^, and 1×10^−12^.

**Figure S5. Regional association plots for new loci.** Results are plotted using the cleft subtype for with the smallest p-values in the genome-wide meta-analyses. Plots were generated using LocusZoom (Pruim et al., 2010). Symbols are colored by linkage disequilibrium in European populations (1000 Genomes Nov. 2014 release).

**Figure S6. Gene expression analysis for genes in Cleft Map loci.** For all genes in the topologically associated domains contained Cleft Map SNPs, genes are color-coded based on their craniofacial co-expression module from Hooper et al.

**Table S1. Samples used in meta-analyses**

**Table S2. Results for all SNPs analyzed in the cleft map**

**Table S3. Permutation results for all SNPs analyzed in the cleft map**

**Table S4. Genomic coordinates of hESC topologically associated domains overlapping Cleft Map SNPs**

**Table S5. Expression data for cleft map regions**

**Table S6. Datasets for SysFACE expression analyses**

