## Supplementary figures and images for "A systematic genetic analysis and visualization of phenotypic heterogeneity among orofacial cleft GWAS signals"

### Figures S1-S6

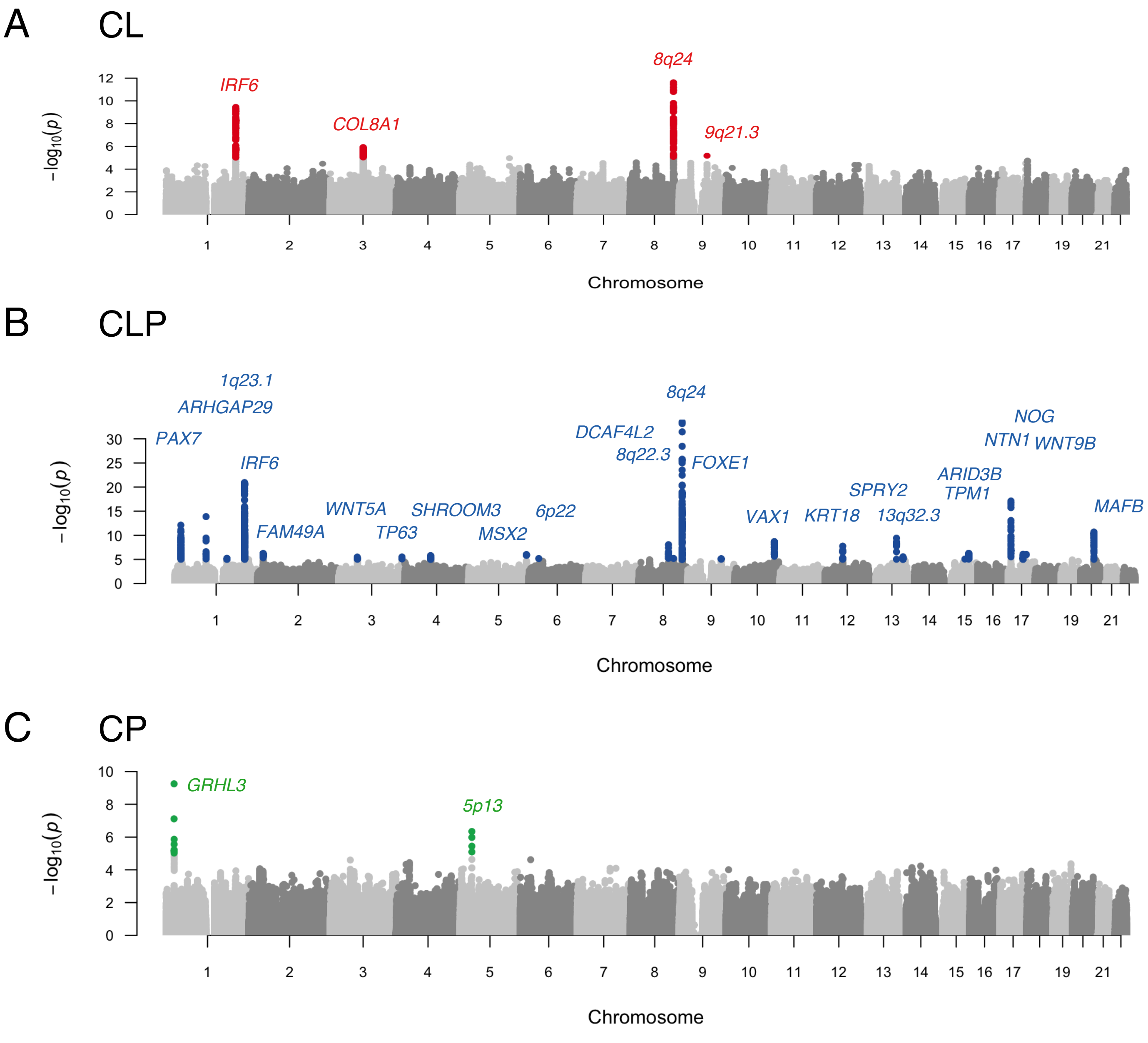

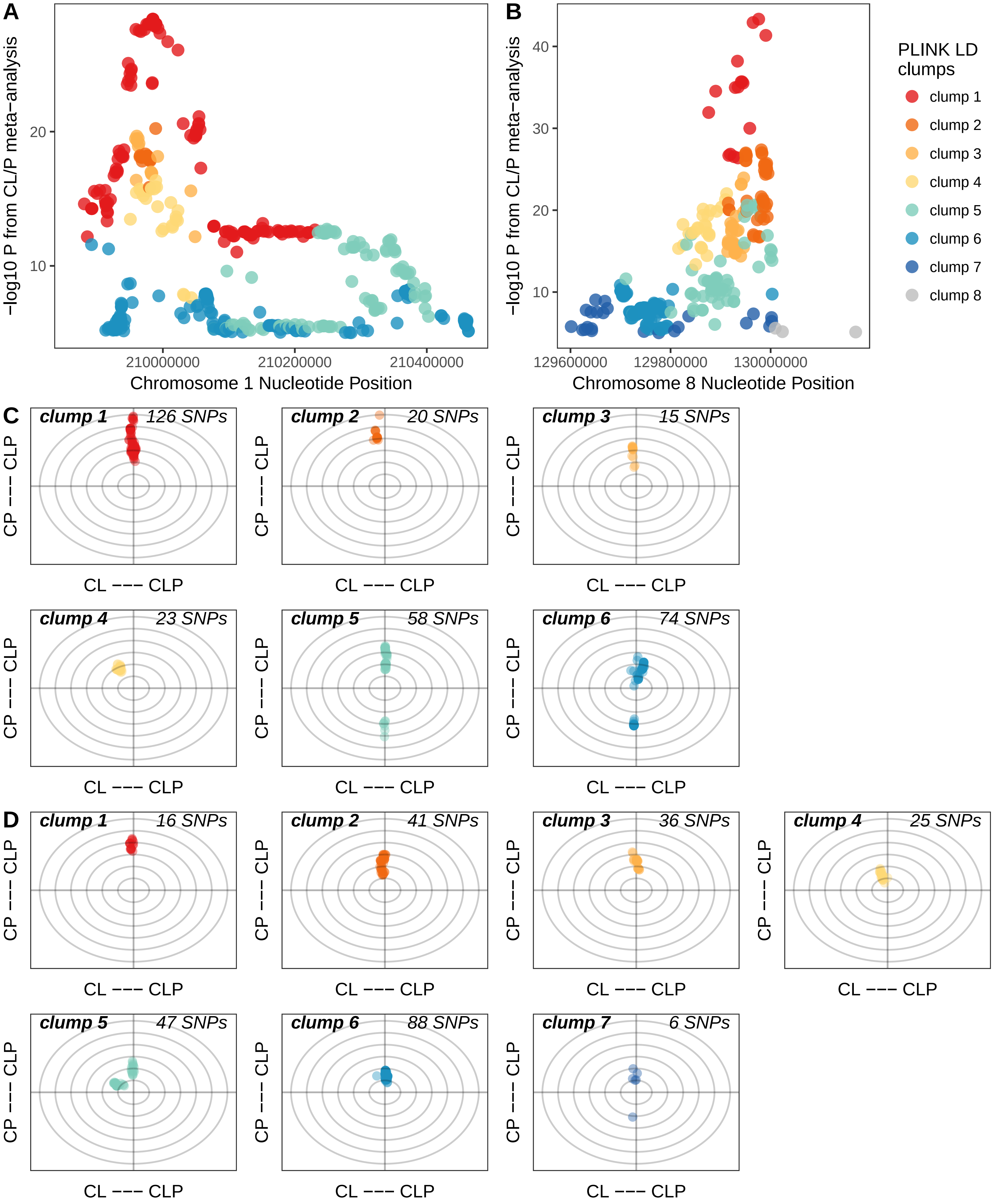

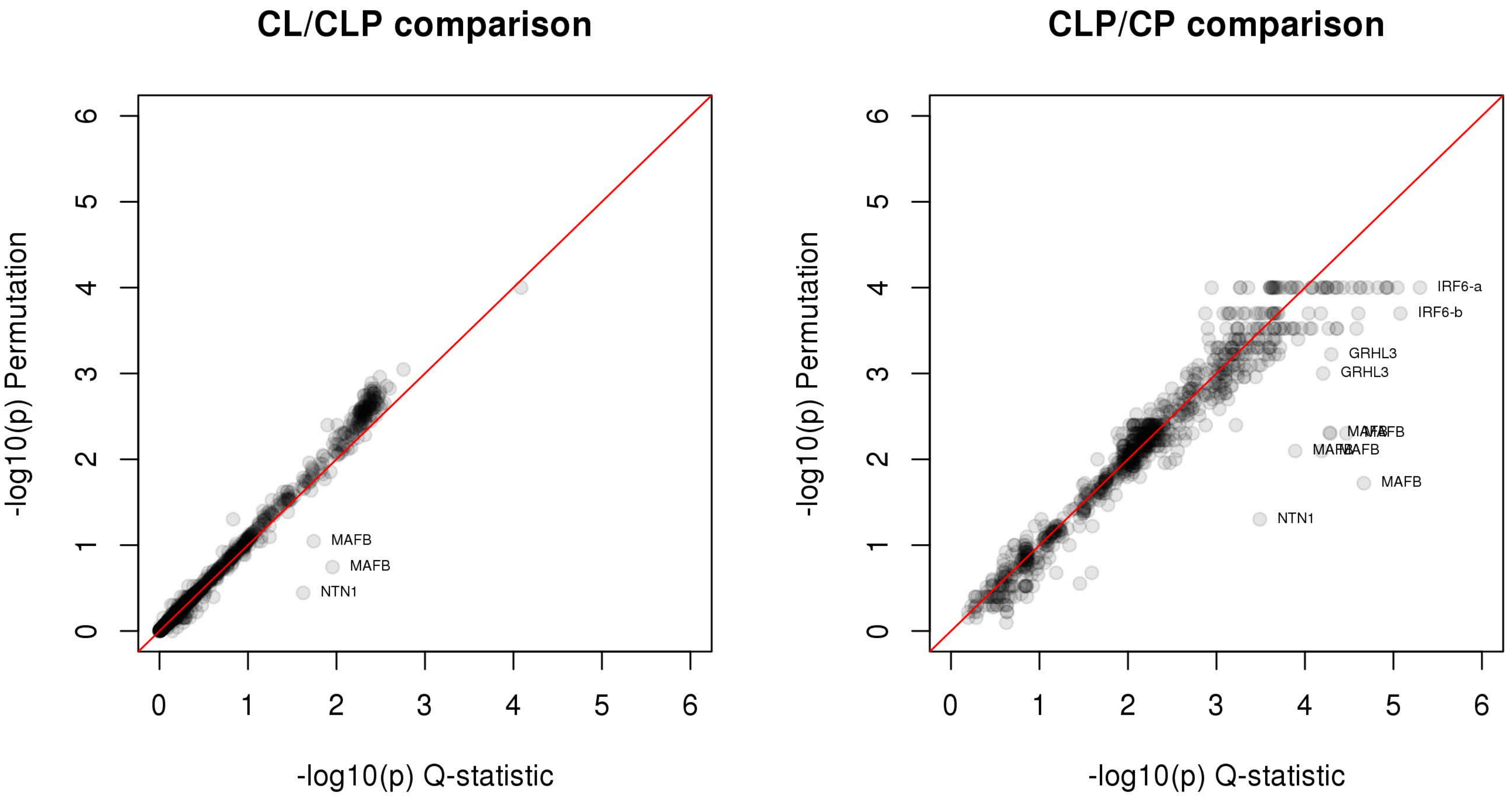

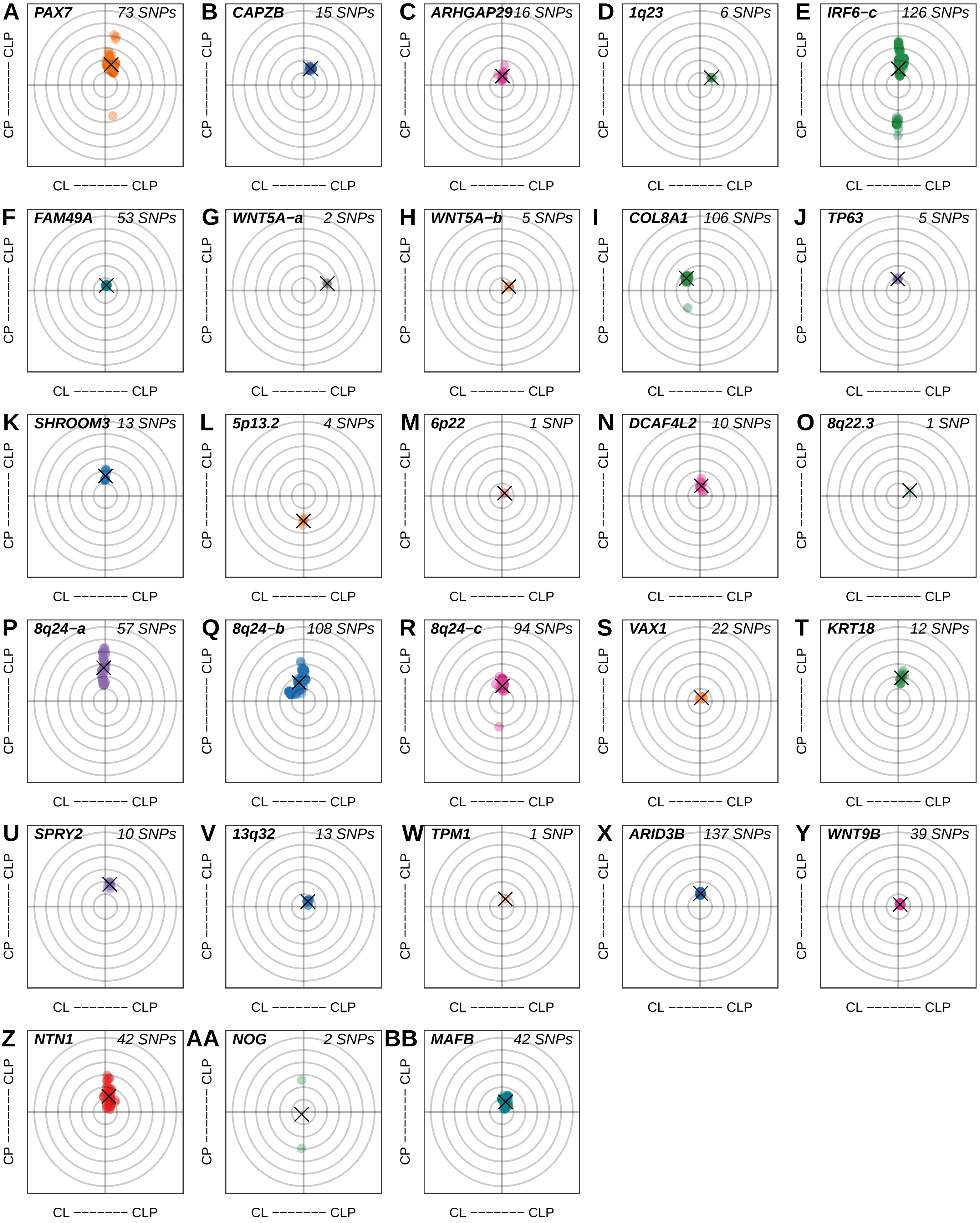

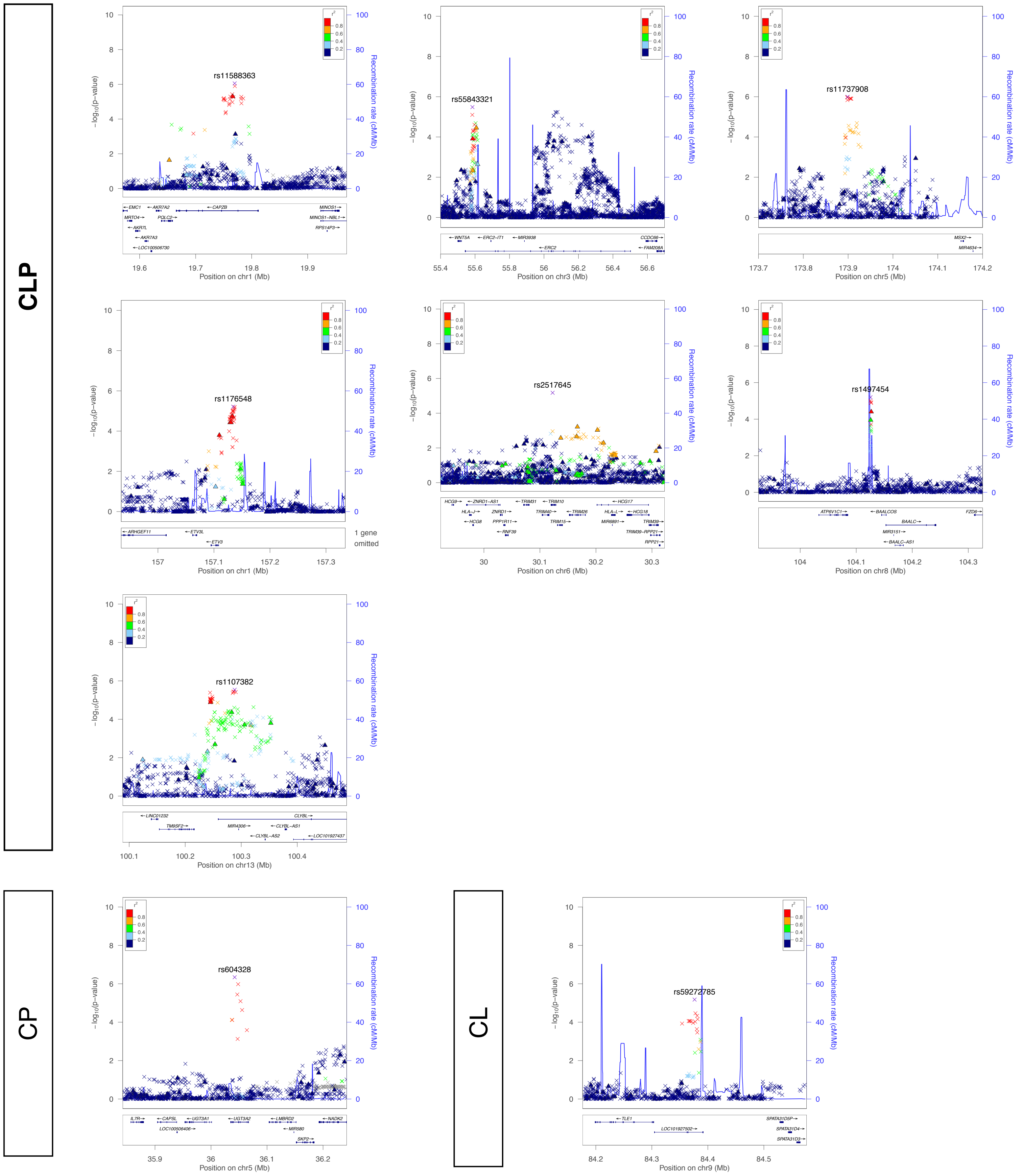

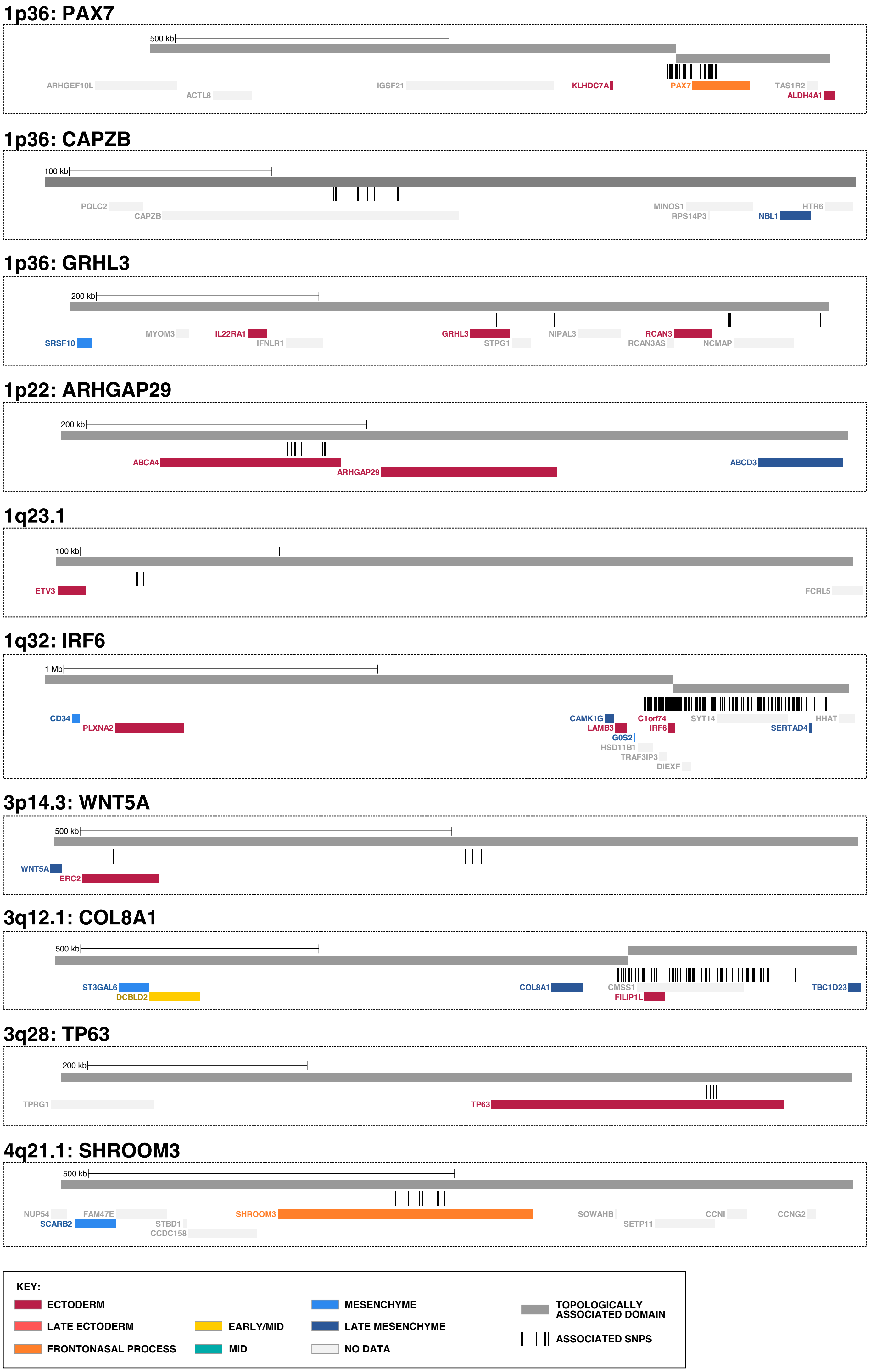

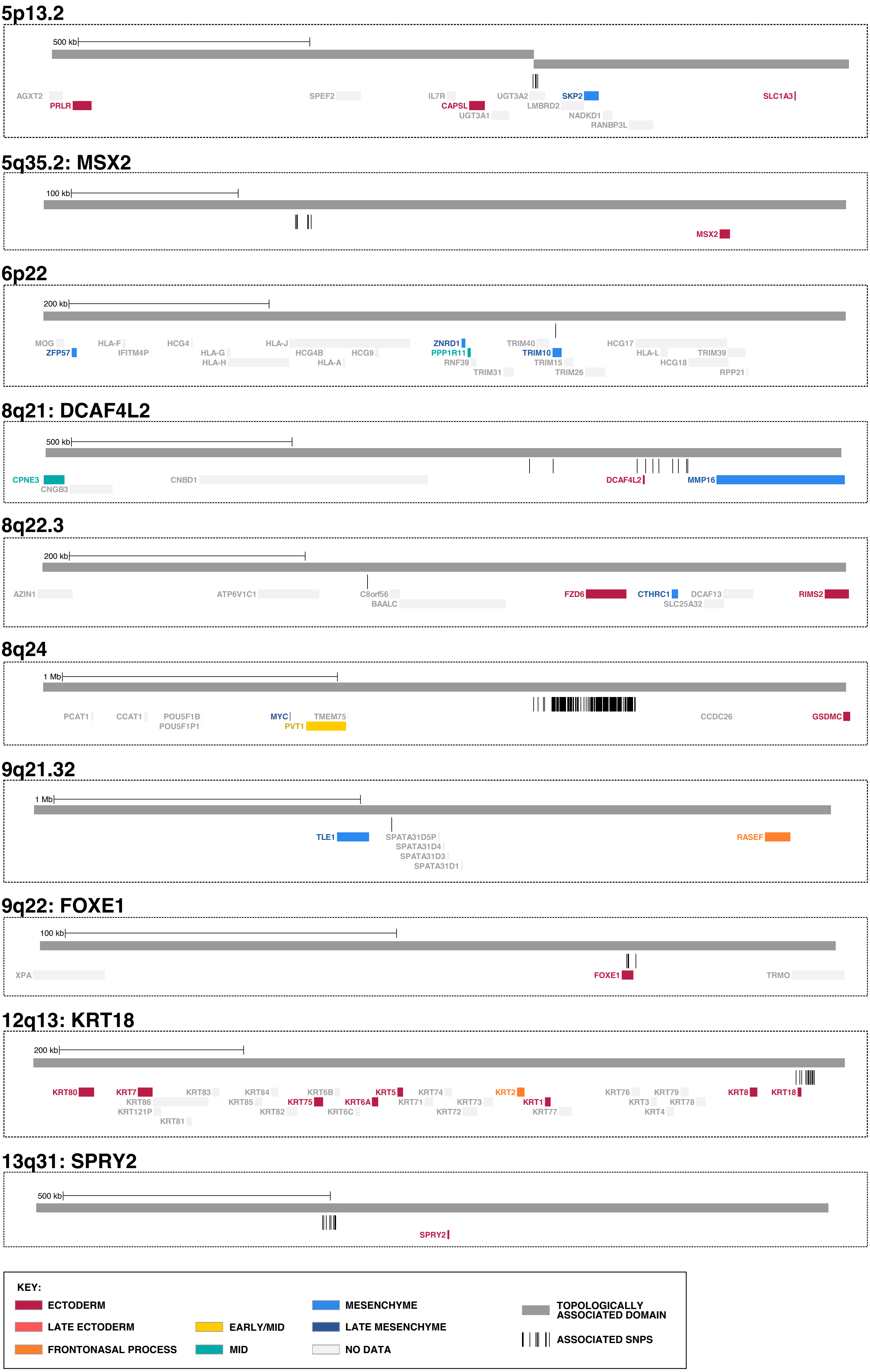

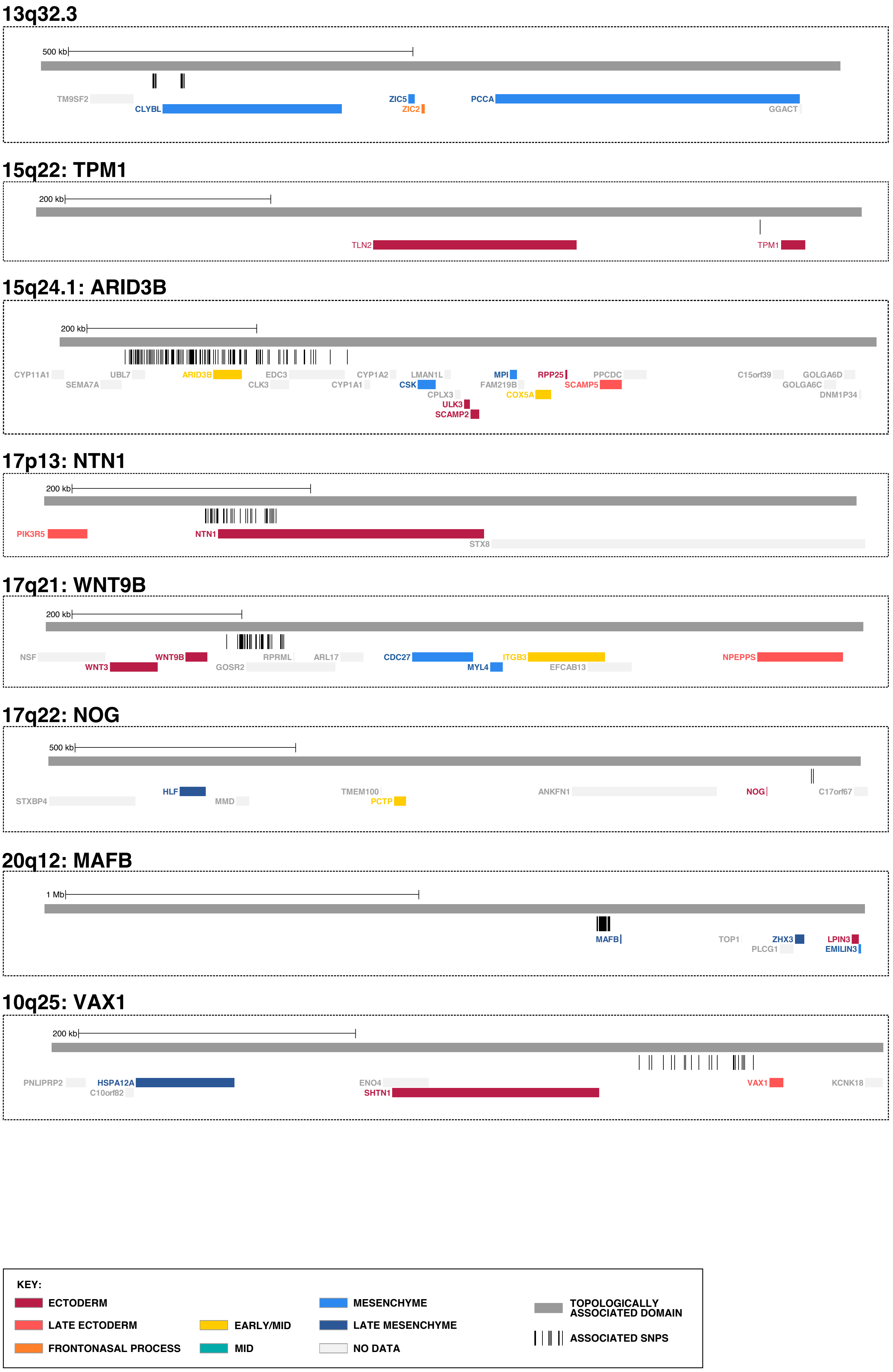
