## Supplementary material for "A systematic genetic analysis and visualization of phenotypic heterogeneity among orofacial cleft GWAS signals": Table S1

| **Table S1. Samples used in meta-analyses** | | | | |
| --- | --- | --- | --- | --- |
| **Phenotype** | **Sample Type** | **POFC** | **GENEVA** | **TOTAL** |
| CL | Trios | 271 | 461 | 732 |
|  | Cases | 179 | -- | 179 |
| CLP | Trios | 1048 | 1143 | 2191 |
|  | Cases | 644 | -- | 644 |
| CP | Trios | 165 | 451 | 616 |
|  | Cases | 78 | -- | 78 |
| Controls |  | 1700 |  |  |
