## Supplementary material for "A systematic genetic analysis and visualization of phenotypic heterogeneity among orofacial cleft GWAS signals": Table S4

| **Table S3. Genomic coordinates of hESC topologically associated domains overlapping Cleft Map SNPs** | |
| --- | --- |
| **Locus** | **Coordinates (hg19)** |
| PAX7 | chr1:17,777,013-19,257,400 |
| IRF6 | chr1:207,951,987-211,500,640 |
| CAPZB | chr1:19,607,414-20,007,413 |
| GRHL3 | chr1:24,264,250-24,991,845 |
| ARHGAP29 | chr1:94,380,850-94,958,350 |
| 1q23 | chr1:157,091,368-157,498,510 |
| FAM49A | N/A |
| WNT5A | chr3:55,499,536-56,589,179 |
| COL8A1 | chr3:98,305,592-100,005,145 |
| TP63 | chr3:188,947,865-189,686,748 |
| SHROOM3 | chr4:77,047,115-78,153,633 |
| 5p13.2 | chr5:34,987,573-36,738,166 |
| MSX2 | chr5:173,745,519-174,230,877 |
| 6p22 | chr6:29,607,111-30,416,039 |
| DCAF4L2 | chr8:87,519,835-89,355,996 |
| 8q22.3 | chr8:103,846,650-104,533,481 |
| 8q24 | chr8:127,836,153-130,785,484 |
| 9q21.32 | chr9:83,185,515-85,831,944 |
| FOXE1 | chr9:100,437,903-100,682,724 |
| VAX1 | chr10:118,362,310-118,979,720 |
| KRT18 | chr12:52,513,734-53,393,733 |
| SPRY2 | chr13:80,126,308-81,642,647 |
| 13q32.3 | chr13:100,058,049-101,264,656 |
| TPM1 | chr15:62,609,582-63,416,074 |
| ARID3B | chr15:74,643,840-75,620,983 |
| NTN1 | chr17:8,770,548-9,466,485 |
| WNT9B | chr17:44,755,711-45,731,967 |
| NOG | chr17:53,017,278-54,900,403 |
| MAFB | chr20:37,672,345-40,023,417 |
