## Supplementary material for "A systematic genetic analysis and visualization of phenotypic heterogeneity among orofacial cleft GWAS signals": Table S6

| **Table S5. SysFACE Gene expression datasets** | | | |
| --- | --- | --- | --- |
| **Tissue** | **Developmental Time Points** | **Microarray** | **RNA-Seq** |
| Platform |  | Affymetrix GeneChip Mouse Genome 430 2.0 Array | Illumina HiSeq2500 |
| Mandibular Process | E10.0, E10.5, E11.0, E11.5, E12.0, E12.5 | GEO: GSE7759 |  |
| Maxillary Process | E10.5, E11.0, E11.5, E12.0, E12.5 | GEO: GSE7759 |  |
| Frontonasal Prominence | E10.5, E11.0, E11.5, E12.0, E12.5 | GEO: GSE7759 |  |
| Whole Body Reference |  | GEO: GSE32334 | unpublished |
| Palate | E13.5 | FaceBase: FB00000468.01 |  |
| Palate | E14.5 | FaceBase: FB00000474.01  GEO: GSE11400 | FaceBase: FB00000768.01 (posterior)  FaceBase: FB00000769.01 (anterior) |
| Palate | P0 | GEO: GSE31004 |  |

NCBI GEO: <https://www.ncbi.nlm.nih.gov/geo/>

FaceBase: <https://www.facebase.org>
